# Cognitive enrichment preserves retrosplenial parvalbumin density and cognitive function in female 5xFAD mice

**DOI:** 10.1101/2025.01.15.633249

**Authors:** Dylan J. Terstege, Jonathan R. Epp

## Abstract

**BACKGROUND:** The rate of cognitive decline in Alzheimer’s disease (AD) varies considerably from person to person. Numerous epidemiological studies point to the protective effects of cognitive, social, and physical enrichment as potential mediators of cognitive decline in AD; however, there is much debate as to the mechanism underlying these effects. The retrosplenial cortex (RSC) is one of the earliest brain regions with impaired functions during AD pathogenesis, and its activity is affected by cognitive, social, and physical stimulation.

**METHODS:** In the current study, we use the 5xFAD mouse mode of AD to examine the impact of enriched housing conditions on cognitive function in AD and the viability of a particularly vulnerable cell population within the RSC – parvalbumin interneurons (PV-INs).

**RESULTS:** Enriched housing conditions improved cognitive performance in 5xFAD mice. These changes in cognitive performance coincided with restored functional connectivity of the RSC and preserved PV-IN density within this region. Along with preserved PV-IN density, there was an increase in the density of perineuronal nets (PNNs) across the RSC of 5xFAD mice housed in enriched conditions. Direct manipulation of PNNs revealed that these extracellular matrix structures protect PV-INs from amyloid toxicity.

**CONCLUSIONS:** Together, these results provide support for the PNN-mediated maintenance of PV-INs in the RSC as a potential mechanism mediating the protective effects of enrichment against cognitive decline in AD.

## BACKGROUND

Alzheimer’s disease (AD) is a neurodegenerative disorder characterized by progressive cognitive decline. Along with cognitive changes, AD is associated with pathological changes including an accumulation of amyloid plaques and neurofibrillary tangles [1]. However, changes in brain function can be detected even prior to the development of these pathological signs [2,3]. Furthermore, the degree of amyloid plaque and neurofibrillary tangle accumulation are relatively poor predictors of t cognitive decline [4–6]. One factor that complictes the predictive power of these hallmark pathological signs is environmental enrichment.Individuals with a high degree of cognitive, social, and physical stimulation show an increased resilience to cognitive decline even when faced with similar pathological burden [7–9].

Numerous epidemiological studies have identified protective effects of cognitive, social, and physical enrichment against disease progression in AD [10–12]. However, the underlying mechanisms which promote this cognitive resilience are not yet fully understood. Many studies have focused on isolating the individual influences of cognitive, social, and physical enrichment on resilience to cognitive decline. While this has great value in identifying potential specific mechanisms which may underlie the protective effects of each specific intervention; it has been demonstrated that these manipulations are most effective in tandem [13]. This combined enrichment likely gives rise to a wide variety of enhancements throughout the brain. In the current study, we have focused on the interplay between enrichment and retrosplenial cortex (RSC) dysfunction, early predictor of the transition between MCI and AD [14].

One of the earliest functional changes that can be detected in the brains of individuals prior to the onset of AD is dysfunction within the RSC [15–17]. Hypometabolism in the retrosplenial cortex can be detected years before AD diagnosis [18,19] and, the severity of RSC hypometabolism may be predictive of transition from MCI to AD [14,20–22]. Similarly, several reports have identified impaired activity and expression of parvalbumin-expressing interneurons (PV-INs) in AD, particularly in the RSC [23–26]. These fast-spiking interneuron populations are the most abundant source of GABAergic signalling in the RSC and are critical in maintaining the excitatory-inhibitory balance of the region. Due to the fast-spiking nature of their firing, PV-INs have considerably high metabolic demands and are highly vulnerable to hypometabolism.

Here, we examined the effects of long-term multi-modal enrichment on cognitive performance and RSC function. We demonstrate impaired cognitive function in 5xFAD mice housed under standard housing conditions, but not in 5xFAD mice housed in enriched environments. Underlying impaired cognitive performance, 5xFAD mice housed under standard conditions displayed disrupted functional connectivity of the RSC and impaired expression of PV-INs. These impairments were not observed in 5xFAD mice housed in enriched environments. Finally, the density of perineuronal nets (PNNs) in the RSC was impaired in 5xFAD mice housed under standard conditions, with remaining PNNs expression being more closely restricted to PV-INs. Together, these results suggest PV-IN maintenance as a potential mechanism through which enrichment promotes resilience to cognitive decline in AD.

## METHODS

### MICE

Female mice were used for all experiments. For enrichment experiments, heterozygous 5xFAD mice were produced via *in vitro fertilization by the* University of Calgary Centre for Genome Engineering. Sperm was obtained from male 5xFAD mice (#034840-JAX) and female C57Bl/6J mice were used as oocyte donors (#000664-JAX). Mice were 8 weeks old at the start of the experiment and all procedures were complete by the time these mice were 6 months old (24 weeks). For amyloid and chondroitinase infusion experiments, homozygous 8-week-old C57Bl/6J mice were bred from breeders purchased from the Jackson Laboratory (Bar Harbour, ME). For all experiments, the room lighting in housing facilities was maintained on a 12 h/12 h light/dark cycle (8 am, lights on). All procedures were conducted during the light cycle phase. All experiments involving animal subjects were conducted in accordance with Canadian Council on Animal Care guidelines and with the approval of the University of Calgary Animal Care Committee.

### ENRICHMENT

Custom enrichment cages, based on a previous design [27], were constructed to provide continuous enrichment. These large cages measured 61 cm X 40.5 cm X 37.5 cm (length X width X height) and contained a divider which separated a compartment containing food from a second compartment containing water. To pass from one compartment to the other, mice had to climb up a ladder to the second level of these cages and pass through an interchangeable maze insert. Maze inserts served as cognitive enrichment and were changed every 3 days. 6 maze inserts were rotated through each cage and, over time, additional removable walls were added within these mazes to further modify their design. Each maze configuration was used only once, throughout the entire 16-week enrichment protocol and, the configurations were identical each enrichment chamber. In addition to interchangeable maze inserts, mice in the enriched condition were also given access to running wheels for physical enrichment. Additionally, with larger cages mice received social enrichment via an increased number of cage mates (11 – 12 mice per cage). Mice in the control conditions were housed in standard cages with 4 – 5 mice per cage and free access to food and water.

### BEHAVIOURAL TESTING

Mice were habituated to handling for 5 days prior to all behavioural tasks. Mouse behaviour was tracked using ANYmaze Behavioural Tracking Software (Stoelting Co., Wood Dale, IL, USA).

#### Y Maze Spontaneous Alternation

Spatial working memory ability was assessed using a Y maze spontaneous alternation task. The arms of the Y maze apparatus were 41.5 cm long and 7.5 cm wide and was cleaned with 70% EtOH before and after each trial. Mice were placed at the end of one arm of the Y maze allowed 5 min to explore the apparatus. During this trial, the number of times that mice entered each of the three arms of the maze sequentially without re-entering any of the two previously entered arms, it was considered a spontaneous alternation. The number of spontaneous alternations was expressed as a percentage of the total number of arm entries for statistical comparisons between groups.

#### Trace Conditioning

Trace conditioning took place in Ugo Basile (Gemonio, Italy) contextual fear conditioning chambers (17 cm X 17 cm) placed inside of sound attenuating cabinets. The conditioning chamber had black and white vertical stripes along the walls, metal bars along the floor, and was cleaned with 30% isopropyl alcohol before and after each trial. During a conditioning trial, mice were allowed to explore the chamber for 3 min prior to the onset of a 20 s tone (2700 Hz). A 20 s trace period followed the offset of this tone prior to the delivery of a foot shock (1 mA, 2 s). The tone-trace-shock presentation was repeated 4 times (5 presentations in total) with a 200 s intertrial interval. Mice were removed from the conditioning chamber 1 min after the final shock and returned to the home cage.

Trace memory testing took place 24 h after the conditioning trial in a distinct context. These chambers had solid grey walls and floors and an open vial of vanilla extract underneath the floor. Chambers were cleaned with 70% EtOH before and after each trial. During this trial, mice were allowed to explore the chamber for 2 min prior to the onset of a 20 s tone. Tones were presented with an intertrial interval of 220 s, with a total of 3 tones having been presented. During subsequent analyses, the 20 s following the offset of the tone was considered to be the trace period.

Contextual memory testing took place 48 h after the conditioning trial, during which mice were returned to the conditioned context for an 8 min trial without any shocks or tones. In both testing sessions, freezing was used as the primary measure of memory retention and was defined as a complete lack of motion, except for respiration, for at least one second.

### SURGICAL PROCEDURES

Surgeries were conducted under isoflurane anesthesia delivered via a Somnosuite anesthetic delivery system (Kent Scientific). Mice were induced at 5% isoflurane, transferred to a stereotaxic frame, and maintained at 1-2% isoflurane. Mice were administered analgesia (Metacam, 5 mg/kg) and fluid support (warmed saline, 0.5 mL) at the beginning of the procedure and monitored closely throughout. A robotic stereotaxic manipulator paired with stereodrive software (Neurostar) was used to drill burr holes and guide a glass infusion needle into place. Using a Nanoject III infusion system (Drummond Scientific), all mice received an injection of Chondroitinase (ChABC, prepared at 100 units/mL in saline; Sigma Aldrich, C3667-5UN) at an RSC target in the right hemisphere (AP: -2.2; ML: 0.5; DV: 1.0) and a saline injection at an RSC target in the left hemisphere (AP: - 2.2; ML: -0.5; DV: 1.0). At each target, these injections were delivered in five sets of 100 nL pulses at a rate of 10 nL/s with 10 s between pulses. Half of the mice (n = 5) then received bilateral injections of amyloid-β 1-42 (prepared at 1mg/mL and incubated for 3 days at 36 °C prior to use; TOCRIS, 1428; 100 nL/site), while the other half (n = 5) received bilateral saline injections (100 nL/site). After each series of injections, the pipette was left at the target depth for 5 min following the final pulse to allow for diffusion of the solution. Once all injections were delivered, the incision was closed with suture material and mice were removed from the stereotaxic frame and were monitored for one week during their recovery.

### HISTOLOGY

#### Perfusion and Tissue Processing

Ninety minutes following the completion of the trace conditioning task, mice were deeply anesthetized with isoflurane and transcardially perfused with 0.1 M phosphate-buffered saline (PBS) and 4% formaldehyde. Brains were extracted and postfixed in 4% formaldehyde at 4 °C for 24 h. Brains were then submerged in 30% w/v sucrose solution in PBS for two to three days until no longer buoyant for cryoprotection. Brains were sagittally sectioned 40 μm thick on a cryostat (Leica CM 1950, Concord, ON, Canada) in 12 series. Sections were kept in a buffered antifreeze solution containing 30% ethylene glycol and 20% glycerol in 0.1 M PBS and stored at -20 °C.

#### Immunofluorescent Staining

For all immunofluorescent staining, free-floating tissue sections were washed three times (10 min per was) in 0.1 M PBS at room temperature. Sections were then incubated at room temperature in primary antibody solution, containing primary antibody, 3% normal goat serum, and 0.3% Triton X-100 (see Supplementary Table S1). Following primary incubation, tissue was washed in 0.1 M PBS (3 x 10 mins) prior to incubation in secondary antibody solution containing secondary antibody and 0.1 M PBS at room temperature for 24 h (see Supplementary Table S1).

PNNs were visualized using Wisteria Floribunda Lectin (WFA) staining[28]. Tissue sections were washed in 0.1 M PBS (3 x 10 mins) prior to incubation in carbo-free blocking buffer (VectorLABS) with 0.2% triton-x100 for 30 mins. Then, sections were stained in a 1:100 dilution of TRITC labeled WFA (VectorLABS) in carbo-free blocking buffer with 0.05% tween-20 for 24 h.

Finally, all tissues were counterstained with DAPI (20 mins, 1:1000 in 0.1 M PBS) before being washed in 0.1 M PBS (2 x 10 mins) and mounted to glass slides. Slides were coverslipped with PVA-DABCO mounting medium.

### IMAGING AND IMAGE ANALYSIS

#### Density Analyses

The density of parvalbumin, c-Fos, and amyloid expression in mouse brain tissue was assessed using images collected at 10X magnification (N.A. 0.4) using an OLYMPUS VS120-L100-W slide scanning microscope (Richmond Hill, ON, CA). Labels were segmented from background based on label size and fluorescent intensity using the user-guided machine learning image processing software *Ilastik*. Binary segmented labels were exported from *Ilastik* and their expression density was assessed within the RSC based on a tracing of this region in the accompanying DAPI channel image in ImageJ.

The expression density of synaptotagmin-2 across the RSC was assessed using images collected at 60X magnification using an oil immersion objective (N.A. 1.42) equipped to an OLYMPUS FV3000 confocal microscope (Richmond Hill, ON, CA). Volumetric imaging frames (10 z steps; 0.35 um z spacing) were collected from the RSC and synaptotagmin-2 puncta were segmented via *Ilastik* from maximum intensity projections of these stacks. The density of these labels were assessed using ImageJ.

#### Functional Connectivity Analyses

Brain-wide c-Fos expression was imaged using an OLMYPUS VA120-L100-W slide scanning microscope (Richmond Hill, ON, CA) equipped with a 10X objective (N.A. 0.4). Using *Ilastik*[29], c-Fos+ cells were segmented to generate binary masks of brain-wide c-Fos expression. These binary masks were then registered to the Allen Mouse Brain Reference Atlas using a modified protocol building upon *Whole Brain*[30,31]. Using custom MATLAB analyses, outputs were organized to yield c-Fos expression densities across 50 neuroanatomical regions (for a detailed list of regions, see Supplementary Table S2). Regional c-Fos expression density was correlated across groups for all possible combinations of regions, generating correlation matrices of regional co-activation for each group[32–34].

### STATISTICS AND DATA VISUALIZATION

All statistical analyses were performed in Prism (GraphPad Software, Version 9.4.0) or with standard MATLAB functions and statistical tests. Independent t-tests, two-way ANOVAs, and Pearson’s Correlations were performed. Data in graphs is presented as mean ± SEM Hypothesis testing was complemented by estimation statistics for each comparison using estimationstats.com[35]. For these estimation statistics, the effect size (Cohen’s d) was calculated using a bootstrap sampling distribution with 5000 resamples along with a 95% confidence interval (CI; bias-corrected and accelerated). Data distribution was assumed to be normal, but this was not formally tested. Statistical methods were not used to predetermine study sizes but were based on similar experiments previously published. Experimenters were blinded to the genotype and housing condition of the animals during all analyses. All statistical comparisons and outputs are included in Supplementary Table S3. All plots were generated in Prism or MATLAB. Circle plots were generated using the circularGraph MATLAB function.

## RESULTS

### Female 5xFAD mice housed under enriched conditions do not show cognitive impairments

The 5xFAD mouse model of AD has awell-characterized timeline of pathological progression and cognitive impairments. At 6 months of age, we observed the expected high density of amyloid-β plaque deposition in the RSC of 5xFAD mice. Enrichment did not influence the accumulation of amyloid-B plaques compared to standard housing (Supplementary Figure 1B). Based on previous characterization of the 5xFAD mice we also expected that deficits in spatial and conditioned memory should arise by 6 months of age. Spatial memory performance assessed in the Y maze spontaneous alternation task supported these previous findings. Here, no effects of genotype or housing condition were observed in 4-month-old mice (Figure 1C). However, by 6 months we observed a lower proportion of spontaneous alternations in 5xFAD mice, and an increase in spontaneous alternations in mice housed under enriched conditions (Figure 1D). Subsequent analyses of the effect sizes of these changes by housing condition revealed that the performance of 5xFAD mice was only impaired relative to WT controls when housed under standard conditions.

**FIGURE 1.**
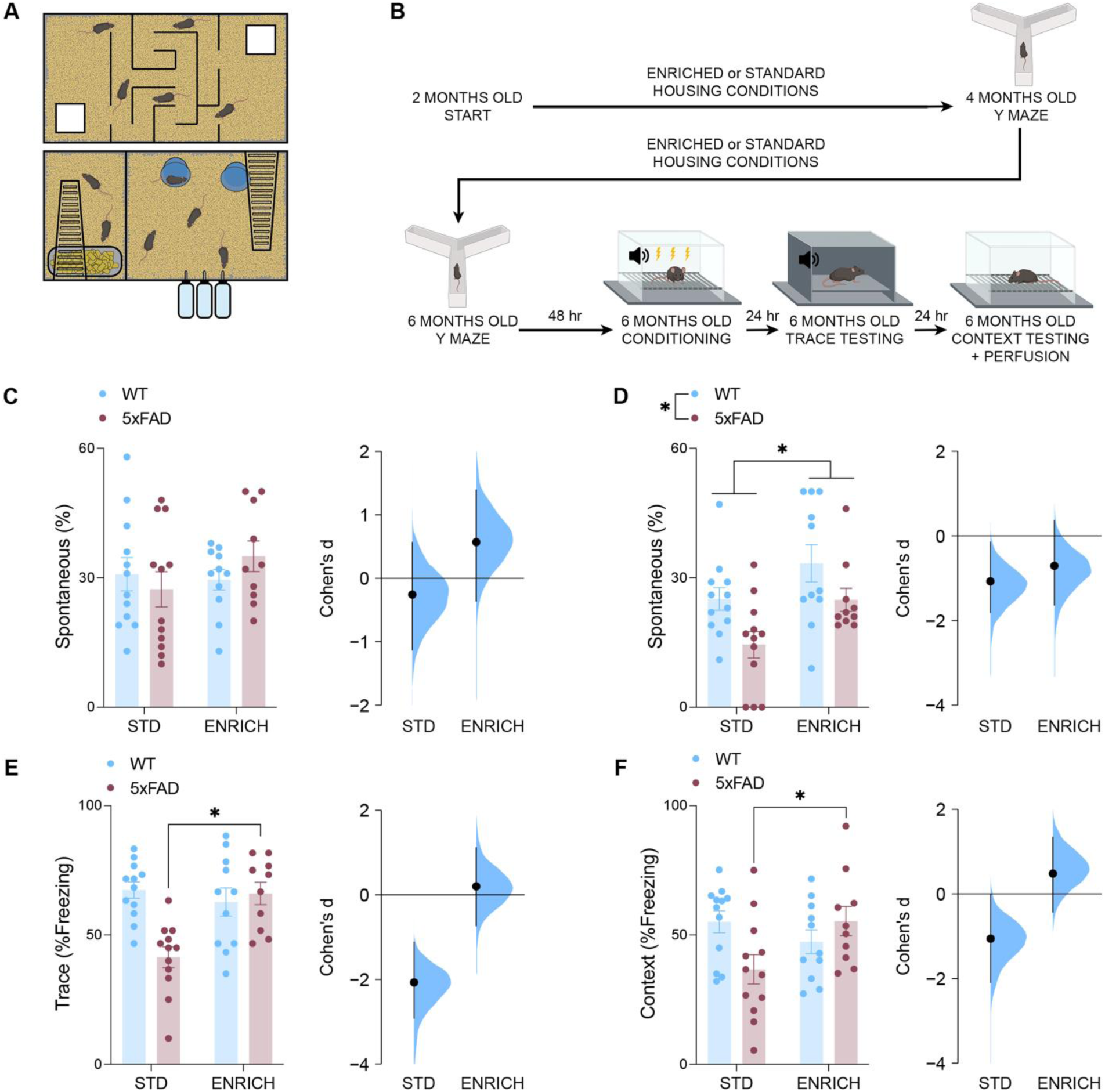
**A** Custom-built multi-level cage designed to provide enrichment through cognitive, social, and exercise-related means. **B** Timeline of behavioural testing and enrichment. **C** Percentage of arm entries in the Y maze which were spontaneous alternations at 4-months-old. Effect size between WT and 5xFAD mice across control (Cohen’s D = -0.255) and enriched (Cohen’s D = 0.203) conditions. **D** By 6-months-old, 5xFAD mice perform a lower percentage of spontaneous alternations, while mice housed in enriched conditions perform a higher percentage of spontaneous alternations (two-way ANOVA, effect of genotype, *F*_(41)_ = 8.606, *P* = 0.0055; effect of housing, *F*_(41)_ = 8.278, *P* = 0.0063). Effect size between WT and 5xFAD mice across control (Cohen’s D = -1.07) and enriched (Cohen’s D = -0.71) conditions. **E** 5xFAD mice housed under standard conditions spent less time freezing during the trace segment of the testing trial (two-way ANOVA, Genotype X Housing interaction, *F*_(41)_ = 11.65, *P* = 0.0015, Šídák’s test, STD 5xFAD vs. ENRICH 5xFAD, *P* = 0.0005). Effect size between WT and 5xFAD mice across control (Cohen’s D = -2.07) and enriched (Cohen’s D = 0.203) conditions. **F** 5xFAD mice housed under standard conditions spent less time freezing upon reintroduction to the conditioned context (two-way ANOVA, Genotype X Housing interaction, *F*_(41)_ = 6.737, *P* = 0.0130, Šídák’s test, STD 5xFAD vs. ENRICH 5xFAD, *P* = 0.0281). Effect size between WT and 5xFAD mice across control (Cohen’s D = -1.06) and enriched (Cohen’s D = 0.482) conditions. Data represent mean ± SEM. *\*P* < 0.05. All statistical comparisons have been provided as Table S3.

Following spatial memory testing, mice were trained on a trace contextual fear memory protocol. This protocol allows for the assessment of both cued and contextual memory. No differences were observed in the rate of freezing behaviour during the initial conditioning trial (Supplementary Figure 1C,D). Following tone presentation during a trace testing trial 24 h later, 5xFAD mice housed under control conditions displayed decreased freezing, indicating impaired retrieval of the cued memory (Figure 1E). Subsequent reintroduction to the original conditioned context revealed that these impairments extended to contextual memory performance, with 5xFAD mice housed under standard conditions exhibiting decreased freezing during this trial (Figure 1F). In each of these trials, 5xFAD mice housed under enriched conditions displayed no memory impairments.

### Enriched housing conditions are sufficient to maintain healthy local activity and global functional connectivity of the retrosplenial cortex in female 5xFAD mice

To investigate the patterns of brain-wide neuronal activity underlying contextual memory retrieval, mice were perfused 90-minutes after reintroduction to a contextually conditioned context. Subsequent c-fos staining revealed peak neuronal activity during contextual memory retrieval. Within the RSC, c-fos expression density was elevated with both standard housing conditions and 5xFAD genotype (Figure 2B). The effect size of this RSC hyperactivity was greatest in 5xFAD mice housed under standard conditions.

**FIGURE 2.**
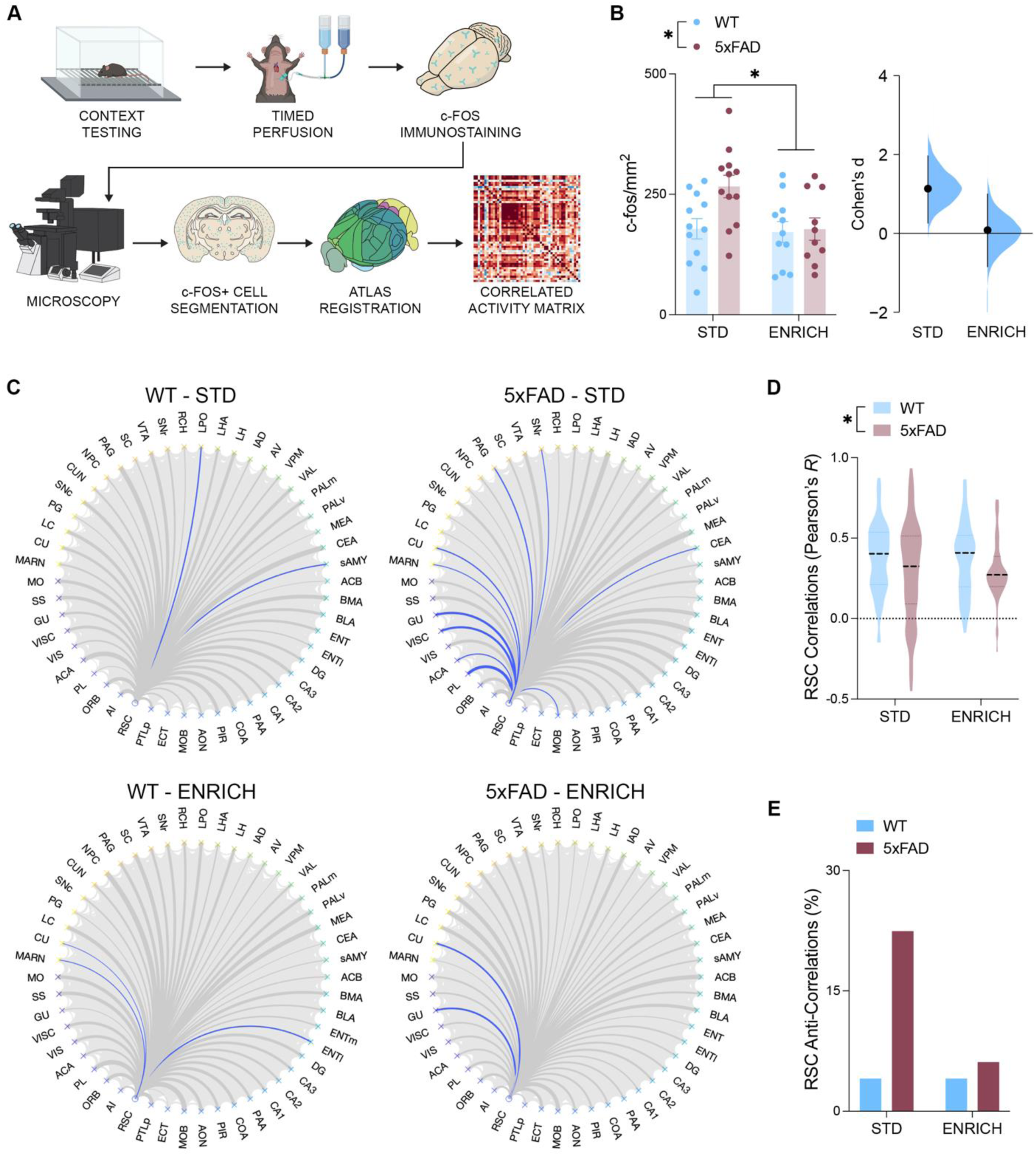
**A** Schematic outlining the process of generating functional connectivity networks from brain-wide c-fos expression. **B** RSC c-fos density was elevated with 5xFAD genotype and standard housing conditions (two-way ANOVA, effect of genotype, *F*_(41)_ = 4.352, *P* = 0.0432; effect of housing, *F*_(41)_ = 4.489, *P* = 0.0402). Effect size between WT and 5xFAD mice across control (Cohen’s D = 1.14) and enriched (Cohen’s D = 0.084) conditions. **C** Circle plots outlining the functional connectome of the RSC. Correlations with positive Pearson’s *r* correlation coefficients are depicted in grey, while correlations with negative correlation coefficients (anti-correlations) are depicted in blue. **D** Correlations involving the RSC were of a lower mean Pearson’s *r* correlation coefficient in 5xFAD mice (two-way ANOVA, effect of genotype, *F*_(192)_ = 5.388, *P* = 0.0219). **E** 5xFAD mice housed under standard conditions (STD) had an increased percentage of anti-correlations relative to all other groups (two-proportion z test, STD WT vs. STD 5xFAD, z = -2.6802, *P* = 0.00736, STD 5xFAD vs. ENRICH WT, z = 2.6802, *P* = 0.00736, STD 5xFAD vs. ENRICH 5xFAD, z = 2.3094, *P* = 0.2088). Data represent mean ± SEM. *\*P* < 0.05. All statistical comparisons have been provided as Table S3.

To determine the impact of RSC hyperexcitability on the ability of this region to communicate effectively with other brain regions during the retrieval of a conditioned memory, c-fos expression density was assessed across 49 other brain regions and correlated with that of the RSC. The resulting correlations outlined the nature of the patterns of co-activity between the RSC and the rest of the network (Figure 2C,D). The mean magnitude of the correlation coefficient between the RSC and the other regions of the network was decreased with 5xFAD status (Figure 2E). However, the number of correlations with a negative sign (anti-correlations) involving the RSC was only elevated above control levels in the 5xFAD mice housed under control conditions, with enriched housing conditions attenuating this effect (Figure 2F).

### Density and synaptic connectivity of parvalbumin interneurons in the retrosplenial cortex is preserved in female 5xFAD mice under enriched housing conditions

It has previously been proposed that anti-correlated functional connectivity may be influenced in part by the activity of PV-INs [36–38]. Furthermore, it has also previously been established that PV-IN density is vulnerable in 5xFAD mice. To assess whether the preserved functional connectivity of the RSC in 5xFAD mice housed under enriched conditions was mediated in part through PV-IN, brains were labelled for this protein. A significant genotype X housing condition interaction revealed a reduction in PV-IN expression density across the RSC of 5xFAD mice housed under standard conditions; however, this deficit was not present in 5xFAD mice housed under enriched conditions (Figure 3B).

**FIGURE 3.**
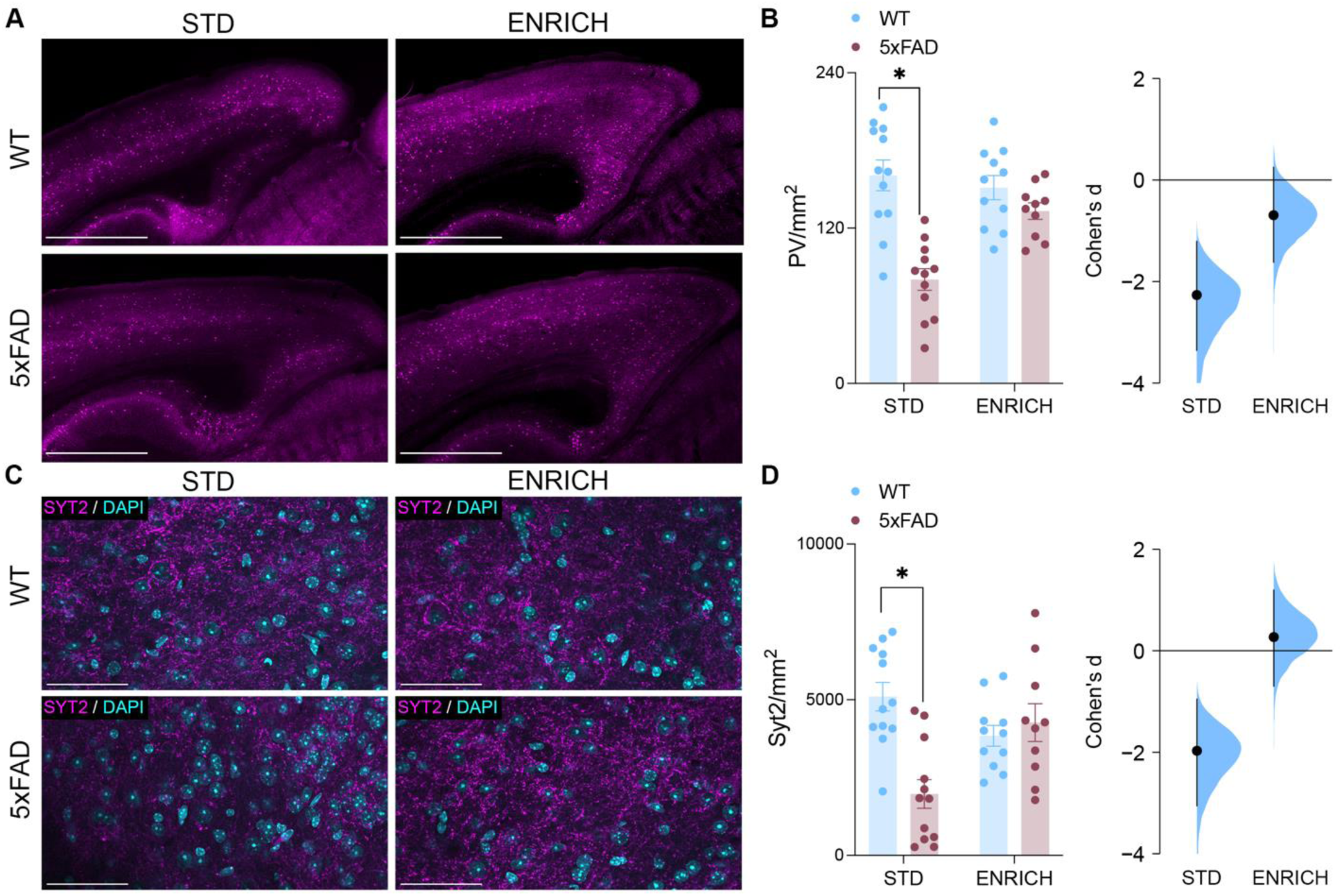
**A** PV staining across the RSC. Scale bars represent 1000 μm. **B** RSC PV density was decreased in 5xFAD mice housed under standard conditions (two-way ANOVA, Genotype X Housing interaction, *F*_(41)_ = 10.83, *P* = 0.0021, Šídák’s test, STD WT vs. STD 5xFAD, *P* < 0.0001). Effect size between WT and 5xFAD mice across control (Cohen’s D = -2.26) and enriched (Cohen’s D = -0.689) conditions. **C** SYT2 staining across the RSC. Scale bars represent 50 μm. **D** RSC SYT2 puncta density was decreased in 5xFAD mice housed under standard conditions (two-way ANOVA, Genotype X Housing interaction, *F*_(41)_ = 14.29, *P* = 0.0005, Šídák’s test, STD WT vs. STD 5xFAD, *P* < 0.0001). Effect size between WT and 5xFAD mice across control (Cohen’s D = -1.97) and enriched (Cohen’s D = 0.275) conditions. Data represent mean ± SEM. *\*P* < 0.05. All statistical comparisons have been provided as Table S3.

To assess the functional consequences of this decrease in PV-IN expression density across the RSC, tissues were stained for synaptotagmin-2 (SYT2). SYT2 is a marker with high specificity for presynaptic contacts from PV-INs; therefore, its expression density provides insight into the ability of PV-INs to regulate both its own activity and the activity of pother cell populations across the region. Here, a significant genotype X housing condition interaction revealed impaired expression density of SYT2 across the RSC of only the 5xFAD mice which were housed under control conditions (Figure 3D).

### Enriched housing conditions are sufficient to preserve the density of perineuronal nets in the retrosplenial cortex of female 5xFAD mice

One factor that has been implicated in the survival of PVINs is the presence of perineuronal nets (PNNs). In the cortex, these extra-cellular matrix structures envelope primarily PV-INs and play important roles in regulating their plasticity while also protecting against oxidative stress. As such, compromised PNN populations may contribute to vulnerability in PV-INs. WFA staining across the RSC revealed a significant genotype X housing condition interaction, wherein 5xFAD mice housed under standard conditions displayed impaired expression density of PNNs across the RSC (Figure 4B). Further histological analyses revealed that, of the PV-INs which remained in the RSC of 5xFAD mice housed under standard conditions, a higher percentage of PV -INs were enveloped by PNNs (Figure 4C). This result alludes to a potentially protective role of PNNs on PV-IN populations in AD neuropathology.

**FIGURE 4.**
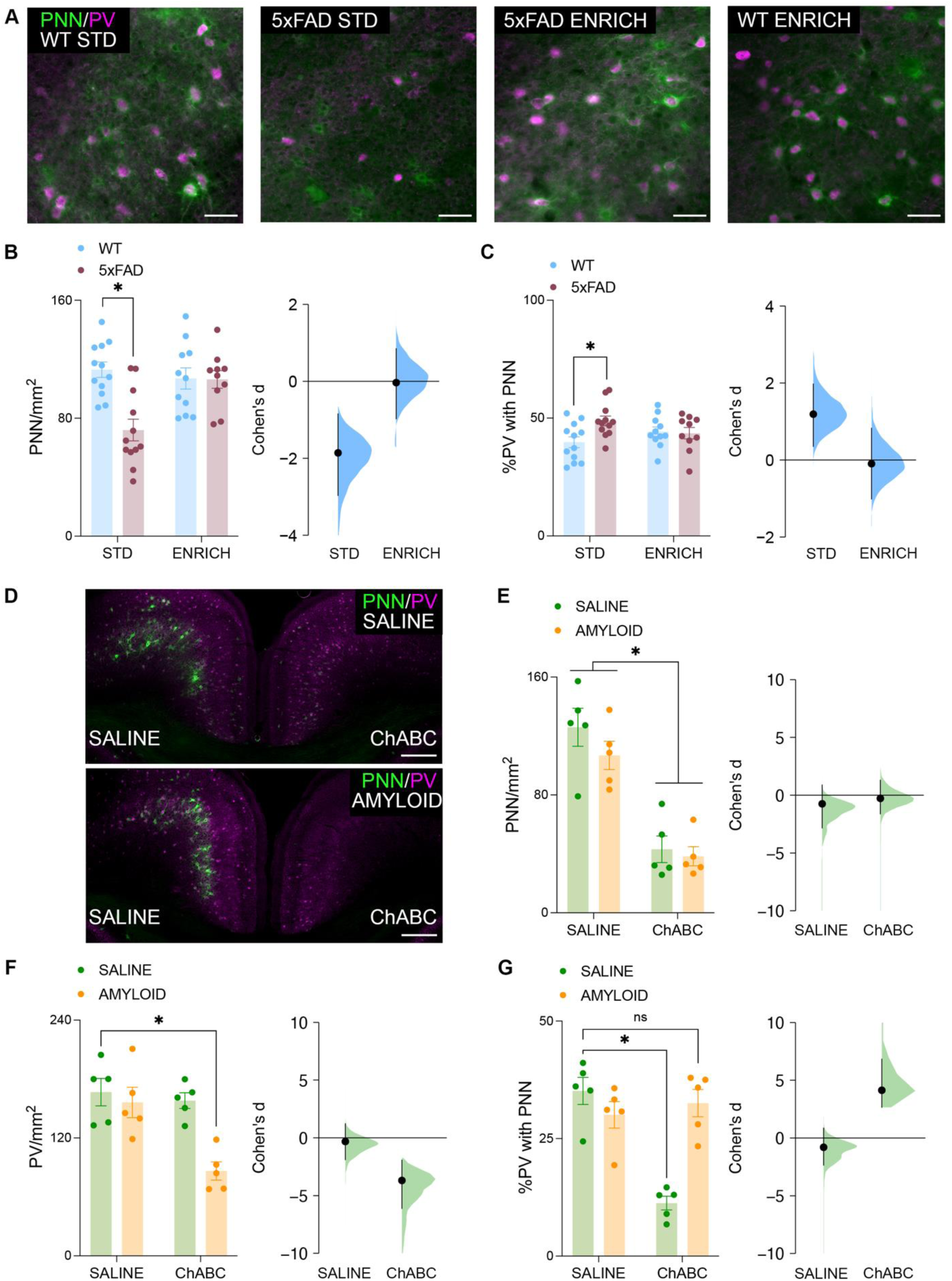
**A** WFA staining for PNNs across the RSC. Scale bars represent 50 μm. **B** RSC PNN density was decreased in 5xFAD mice housed under standard conditions (two-way ANOVA, Genotype X Housing interaction, *F*_(41)_ = 9.498, *P* = 0.0037, Šídák’s test, STD WT vs. STD 5xFAD, *P* < 0.0001). Effect size between WT and 5xFAD mice across control (Cohen’s D = -1.86) and enriched (Cohen’s D = -0.0301) conditions. **C** The percentage of PV cells in the RSC surrounded by PNNs was increased in 5xFAD mice housed under standard conditions (two-way ANOVA, Genotype X Housing interaction, *F*_(41)_ = 4.836, *P* = 0.0336, Šídák’s test, STD WT vs. STD 5xFAD, *P* = 0.0091). Effect size between WT and 5xFAD mice across control (Cohen’s D = 1.19) and enriched (Cohen’s D = -0.0927) conditions. **D** WFA and PV staining across the RSC following bilateral saline (top) or amyloid (bottom) infusion. All mice were also infused with saline (left hemisphere) or ChABC (right hemisphere). Scale bars represent 250 μm. **E** ChABC administration reduced the density of PNNs in the ipsilateral hemisphere relative to the contralateral hemisphere (two-way ANOVA, effect of ChABC, *F*_(16)_ = 60.06, *P* < 0.0001). Effect size between saline- and amyloid-infused mice across control (Cohen’s D = -0.752) and ChABC-infused (Cohen’s D = -0.271) hemispheres. **F** ChABC and amyloid co-administration reduced RSC PV density (two-way ANOVA, Amyloid X ChABC interaction, *F*_(16)_ = 6.368, *P* = 0.0226, Šídák’s test, Saline Saline vs Amyloid ChABC, *P* = 0.0007). Effect size between saline- and amyloid-infused mice across control (Cohen’s D = -0.319) and ChABC-infused (Cohen’s D = -3.69) hemispheres. **G** ChABC administration reduces the percentage of RSC PV surrounded by PNNs, but co-administration of ChABC and amyloid does not (two-way ANOVA, Amyloid X ChABC interaction, *F*_(16)_ = 25.89, *P* = 0.0001, Šídák’s test, Saline Saline vs Saline ChABC, *P* < 0.0001, Saline Saline vs Amyloid ChABC, *P* = 0.4879). Effect size between saline- and amyloid-infused mice across control (Cohen’s D = -0.798) and ChABC-infused (Cohen’s D = 4.14) hemispheres Data represent mean ± SEM. *\*P* < 0.05. All statistical comparisons have been provided as Table S3.

### The presence of PNNs improves the resilience of PV-IN expression to amyloid-β toxicity

In the prior experiment we observed potential protective effect of PNNs on the expression of PV-INs in 5xFAD mice. To further investigate this relationship we used targeted infusions of amyloid-β, ChABC, or the co-administration of bothused As expected, the administration of ChABC reduced the density of PNNs across the RSC of mice perfused 7 days after surgery (Fig. 4D). There was also a significant main effect of amyloid-β on PNN density; however, the magnitude of this effect size was considerably lower than that of ChABC. There was no significant amyloid-β X ChABC interaction effect on the density of PNNs. Conversely, when examining the density of PV-INs, we observed a significant amyloid-β X ChABC interaction (Fig. 4E). Šídák post hoc comparisons further revealed that the density of PV-INs across the RSC was not directly influenced by either treatment on its own but became significantly impaired only when ChABC and amyloid-β were administered together.

Looking more closely at the interaction between surviving PV-INs and PNNs under these conditions, we found that without amyloid-β administration, infusion of ChABC alone caused a reduction in the proportion of PV-INs surrounded by PNNs (Fig. 4F). However, with ChABC and amyloid-β co-infusion, this proportion no longer differs from controls.

## DISCUSSION

Numerous epidemiological studies and previous studies with animal models have identified relationships between different forms of cognitive, social, and physical enrichment and increased resiliency to cognitive decline in AD [39,40]. However, there is little consensus as to the mechanism underlying this effect. As the incidence of AD is expected to keep rising, it is critical that we gain a better understanding of these potentially widely accessible lifestyle interventions which could be integrated into Alzheimer’s disease prevention strategies.

While the underlying mechanisms which contribute most to the protective effects of enrichment on cognitive decline in AD are largely unknown, several neurological correlates have been identified which seem to predict the onset of AD [41–43]. One such correlate is the activity of the RSC, wherein altered activity of this region may predict the onset of cognitive impairment in AD [16,44–46]. The RSC plays important roles across many learning and memory paradigms and it’s activity can be modulated through learning itself, social interactions, and physical exercise [47–52]. Therefore, all three core features of the enrichment protocol in the current study – cognitive, social, and physical enrichment – are expected to influence the activity of this region. The aim of the present study was to examine whether the combination of cognitive, social, and physical enrichment could ameliorate cognitive outcomes and preserve RSC function in the 5xFAD mouse model of AD. With the enrichment protocol beginning prior to the onset of cognitive impairments, the study design mimicked a preventative approach.

Following the conclusion of the enrichment protocol, 5xFAD mice housed in enriched conditions showed improved spatial working memory and contextual memory performance compared to 5xFAD mice housed under standard laboratory conditions. Coinciding with this preservation of cognition, we observed that 5xFAD mice housed under enriched conditions did not display RSC hyperactivity during contextual memory recall like the 5xFAD mice housed under standard conditions did. Furthermore, the density of anticorrelated functional connections formed with the RSC during contextual memory recall was low in 5xFAD mice housed under enriched conditions. This mimicked the density of these connections in the WT mice and was a sharp contrast to the high density of anticorrelated connections observed in 5xFAD mice housed under standard laboratory conditions. These changes in RSC functional connectivity are in alignment with hypotheses which could be derived from previous studies [38]. Increased anticorrelated activity involving this region has previously been reported during early stages of AD, while other reports have shown decreases in RSC anticorrelated activity with physical exercise [53,54]. The present study shows that in AD, the protective effects of enrichment sufficiently suppress RSC dysfunction on a functional connectivity network level.

Based on recent literature demonstrating increased vulnerability of PV-Ins to pathological changes during the early stages of cognitive decline [25,55–57] we examined the impact of enrichment on PV-IN survival. PV-INs are fast-spiking interneurons with high metabolic requirements which have also been thought to contribute to anti-correlated functional connectivity [36,37]. During the early stages of cognitive decline, the RSC becomes hypometabolic which puts considerable strain on PV-INs [21,58,59]. In 5xFAD mice housed under standard conditions, we found a decrease in both the density of PV-INs and PV-IN presynaptic contacts across the RSC. These impairments were not present in 5xFAD mice housed under enriched conditions.

Perineuronal nets are an extracellular matrix structure which have been implicated in both the protection of PV-INs and in regulating their ability to form synaptic connections. In 5xFAD mice housed under standard laboratory conditions, the density of PNNs was reduced across the RSC. Enriched housing conditions were sufficient to keep PNN expression at densities which did not differ from WT groups. PNN depletion has been reported in mouse models of AD as well as post-mortem tissue samples collected from human subjects with AD [60–62]. Furthermore, it has been demonstrated that environmental enrichment is sufficient to preserve PNN density in mouse models of AD [63]. However, we extend beyond these by demonstrating that the PNNs which do remain are more likely to be surrounding PV-INs than other cell types, thereby suggesting that PV-INs themselves may also serve a protective role in the maintenance of PNNs in the RSC of AD models.

To further examine the protective effects of PNNs against AD pathology, PNNs in the RSC were directly manipulated using ChABC. The depletion of PNNs also increased the vulnerability of the PV-INs to this pathology. No significant loss of PV-INs occurred in response to amyloid infusion unless PNNs had been depleted beforehand. Interestingly, the percentage of PV-INs that expressed PNNs was decreased by ChABC treatment as expected however, combined Amyloid and ChABC treatments did not appear to alter this percentage. Considering the loss of PV-INs that we observed following the combined treatment, these results suggest that the PV-INs which survived were those that maintained their PNN expression, further demonstrating the protective role of these structures.

While many studies have reported associations between cognitive, social, and physical enrichment on resilience to cognitive decline in AD; there have been several studies which have been unable to replicate these results [39,64–67]. This has gained relevance in popular culture with mixed results from trials involving commercial “brain-training” applications [68–70]. In presenting our results, it is important to highlight potential confounds which may contribute to the variability in results across the literature. Primarily, one key contributor seems to be the duration of the enrichment protocol [71–73]. Generally, shorter enrichment protocols have yielded smaller effects if any at all [74]. Secondly, the variety of enrichment both within and across modalities is important [75]. Third, early intervention has proven to be most effective in slowing the progression of cognitive decline. Here, we’ve implemented a relatively long multi-modal enrichment protocol prior to the onset of cognitive symptoms to maximize our likelihood of intervention efficacy.

## LIMITATIONS

The current manuscript identifies the protection of PV-INs in the RSC as a mechanism underlying the protective effects of enrichment in AD. Specifically, these results are supported by findings of improved resiliency to amyloid infusion in mice with intact PNNs relative to those which had received ChABC treatment. While this suggests a protective effect of PNNs on PV-IN vulnerability to amyloid toxicity, we acknowledge that amyloid is not the only pathological change which occurs in AD. Future studies should assess the extent to which PNNs are able to protect RSC PV-INs from other pathological changes associated with AD, such as the accumulation of neurofibrillary tau tangles [76,77] and changes in regional glucose metabolism [78,79]. Additionally, it is important to note that the current study was conducted entirely in female mice and as a result it is unknown whether enrichment provides neuroprotective effects in male mice through these same mechanisms.

## CONCLUSIONS

Our findings demonstrate that prolonged cognitive, social, and physical enrichment abolishes cognitive deficits observed in 6-month-old female 5xFAD mice. Our results also suggest that these conditions are sufficient to preserve functional connectivity of the RSC and protect PV-INs in this region, even in the absence of differences in amyloid pathology. Together, these results provide support for the maintenance of PV-INs in the RSC as a potential mechanism mediating the protective effects of enrichment against cognitive decline in AD.

## LIST OF ABBREVIATIONS

AD: Alzheimer’s disease
ChABC: Chondroitinase ABC
CI: Confidence interval
PNN: Perineuronal net
PV-IN: Parvalbumin interneuron
MCI: Mild cognitive impairment
RSC: Retrosplenial cortex
SYT2: Synaptotagmin-2
WT: Wild type

## DECLARATIONS

### ETHICS APPROVAL AND CONSENT TO PARTICIPATE

All experiments involving animal subjects were conducted in accordance with Canadian Council on Animal Care guidelines and with the approval of the University of Calgary Animal Care Committee.

### CONSENT FOR PUBLICATION

Not applicable.

### AVAILABILITY OF DATA AND MATERIALS

Will be made available upon request.

## AUTHOR CONTRIBUTIONS

DJT and JRE conceived and designed the experiments, conducted the analyses, and wrote the manuscript. DJT conducted all experimental procedures. JRE supervised and acquired funding to support all experimental procedures. All authors read and approved the final manuscript.

## ACKNOWLEDGMENTS

We acknowledge the Hotchkiss Brain Institute Advanced Microscopy Platform and the Cumming School of Medicine for support and the use of the Olympus VS120-L100W slide scanning microscope.

