## Supplementary Figure for "Cognitive enrichment preserves retrosplenial parvalbumin density and cognitive function in female 5xFAD mice"

**SUPPLEMENTARY MATERIAL**

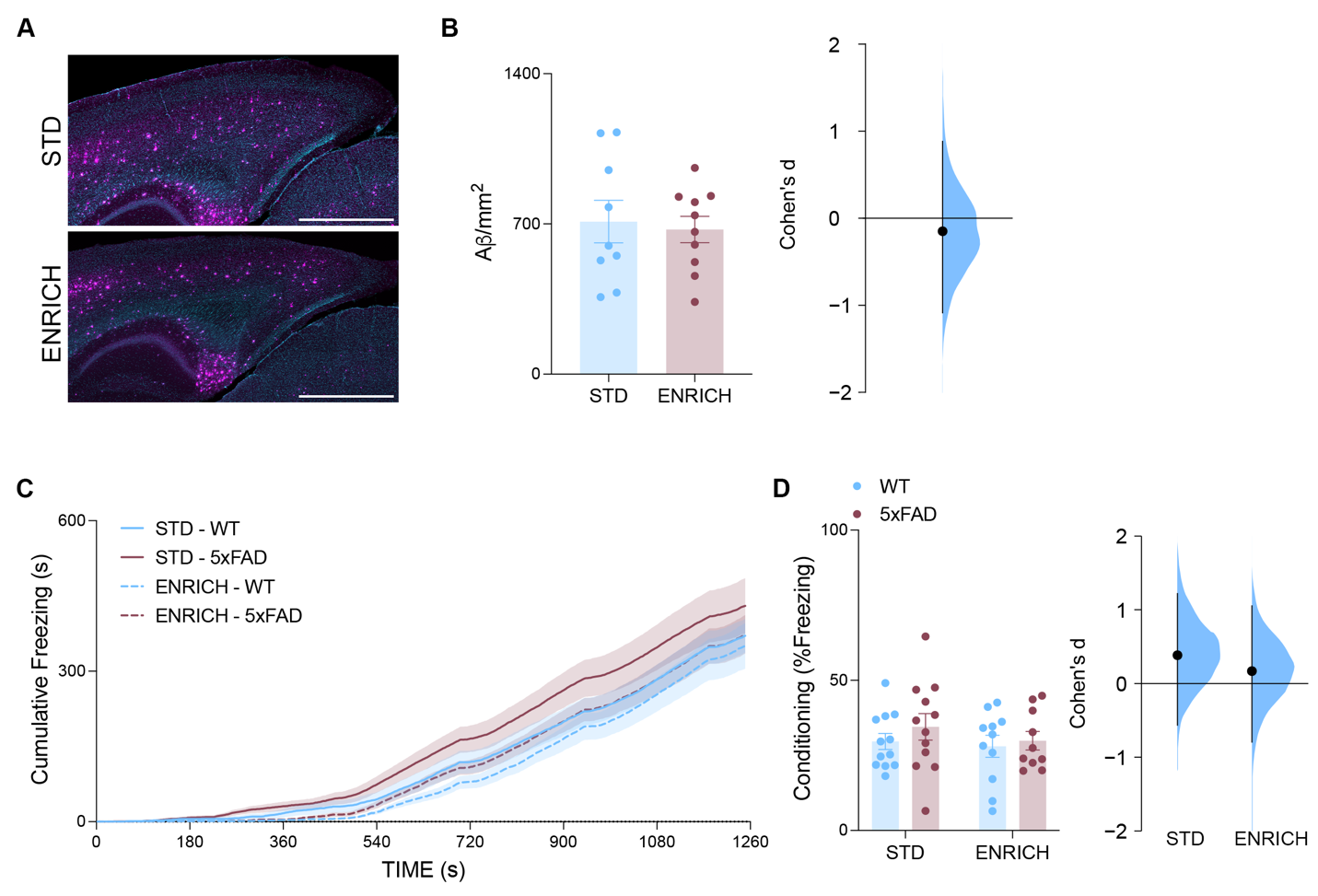

**SUPPLEMENTARY FIGURE 1**

**A** Representative photomicrograph of amyloid-β plaque accumulation across the RSC. Scale bars represent 1000 μm. **B** Density of amyloid-β plaques in the RSC. The density of amyloid-β plaques was not significantly different between 5xFAD mice housed under standard conditions or enriched conditions (two-sample t test, *t*_(17)_ = 0.3210, *P* = 0.7521). Effect size between mice housed in control and enriched conditions (Cohen’s D = 0.745). **C** Cumulative time spent freezing during the conditioning trial of the trace fear conditioning task. **D** Percentage of time spent freezing during the conditioning trial. Effect size between WT and 5xFAD mice across control (Cohen’s D = 0.385) and enriched (Cohen’s D = 0.169) conditions. Data represent mean ± SEM. All statistical comparisons have been provided as Table S3.
